# ShinyCell: Simple and sharable visualisation of single-cell gene expression data

**DOI:** 10.1101/2020.10.25.354100

**Authors:** John F. Ouyang, Uma S. Kamaraj, Elaine Y. Cao, Owen J. L. Rackham

**Affiliations:** Program in Cardiovascular and Metabolic Disorders, Duke-NUS Medical School, Singapore

## Abstract

**Motivation:** As the generation of complex single-cell RNA sequencing datasets becomes more commonplace it is the responsibility of researchers to provide access to these data in a way that can be easily explored and shared. Whilst it is often the case that data is deposited for future bioinformatic analysis many studies do not release their data in a way that is easy to explore by non-computational researchers.

**Results:** In order to help address this we have developed ShinyCell, an R package that converts single-cell RNA sequencing datasets into explorable and shareable interactive interfaces. These interfaces can be easily customised in order to maximise their usability and can be easily uploaded to online platforms to facilitate wider access to published data.

**Availability:** ShinyCell is available at https://github.com/SGDDNB/ShinyCell.

## 1 Introduction

Single-cell RNA sequencing (scRNA-seq) is fast becoming a standard tool for investigating biological phenomena. The resolution that it provides has led to the discovery of novel cell types (See *et al*., 2017), unobserved cell state transitions (Liu *et al*., 2020) or provide the means to perform highly parallel perturbation screens (Replogle *et al*., 2020). In each case, the amount of data that is generated is often far beyond what can be described in a single manuscript and as such it is also increasingly important to share newly generated data in a way that is accessible for both computational and experimental scientists. To this end, we have developed a lightweight R package, named ShinyCell, that can process common scRNA-seq data objects and produce an interactive interface that allows for exploration of the data and easy sharing between collaborators or as part of a publication.

## 2 ShinyCell Workflow and Outputs

Here, we describe the key features of the ShinyCell R package as well as the types of plots that can be generated from a ShinyCell interactive web interface. We then outline basic and advanced workflows to generate these web interfaces.

### 2.1 Key features of ShinyCell

The main purpose of the ShinyCell R package is to provide a simple way to convert existing single-cell data into interactive and shareable web interfaces (Figure 1a). In line with this purpose, we designed ShinyCell with several computational considerations. First, ShinyCell is written in the R programming language (Computing and Others, 2013) and uses the Shiny package (RStudio, 2013) for creating interactive web applications. This allows us to leverage on the visualisation tools in R, in particular the ggplot2 package (Wickham, 2016). Furthermore, the Shiny platform allows for easy sharing on online platforms such as shinyapps.io or be hosted via Shiny Server. Second, ShinyCell uses pre-processed single-cell RNA-seq data in the format of Seurat or SingleCellExperiment (SCE) objects as inputs. The Seurat platform (Butler *et al*., 2018) and Scater pipeline (McCarthy *et al*., 2017) (which uses the SCE object) are two of the most commonly used single-cell analysis tools. More importantly, both tools and ShinyCell are written in R, allowing users who are processing their single-cell data using Seurat or Scater to easily adopt ShinyCell into their existing pipelines. A recent comparison of visualisation tools for scRNA-seq (Cakir *et al*., 2020) highlights that visualisation tools written in the R programming language do not have support for both Seurat or SCE objects while tools that support Seurat or SCE objects are written in other programming languages such as Python. Third, the web interfaces generated by ShinyCell are designed to be memory efficient so that multiple users can access the app simultaneously. The gene expression data is stored using the hdf5 file format on disk and only the expression of genes that are plotted are loaded into memory. We also optimised the chunking in the hdf5 file such that the expression data of each gene is stored contiguously and thus can be accessed quickly. Furthermore, the single-cell metadata is stored in data.table format (Dowle *et al*., 2019) which is faster to manipulate than the core R data.frame format. Fourth, users can export the plots generated in the Shiny app as PDF or PNG images, which can be used for presentation or publication purposes. Fifth, we added support for the inclusion of multiple single-cell datasets into a single Shiny app (see more details in Section 2.4). This is useful to gather multiple datasets into a single interactive interface as an accompanying resource for publication. Sixth, ShinyCell is easy to run and highly expandable. In the simplest form, ShinyCell can convert a Seurat / SCE object into a Shiny app with five lines of code (see more details in Section 2.3). Furthermore, the source code of the generated Shiny app is made transparent to users and users can add additional custom visualisation to the source code if they desire.

**Fig. 1.**
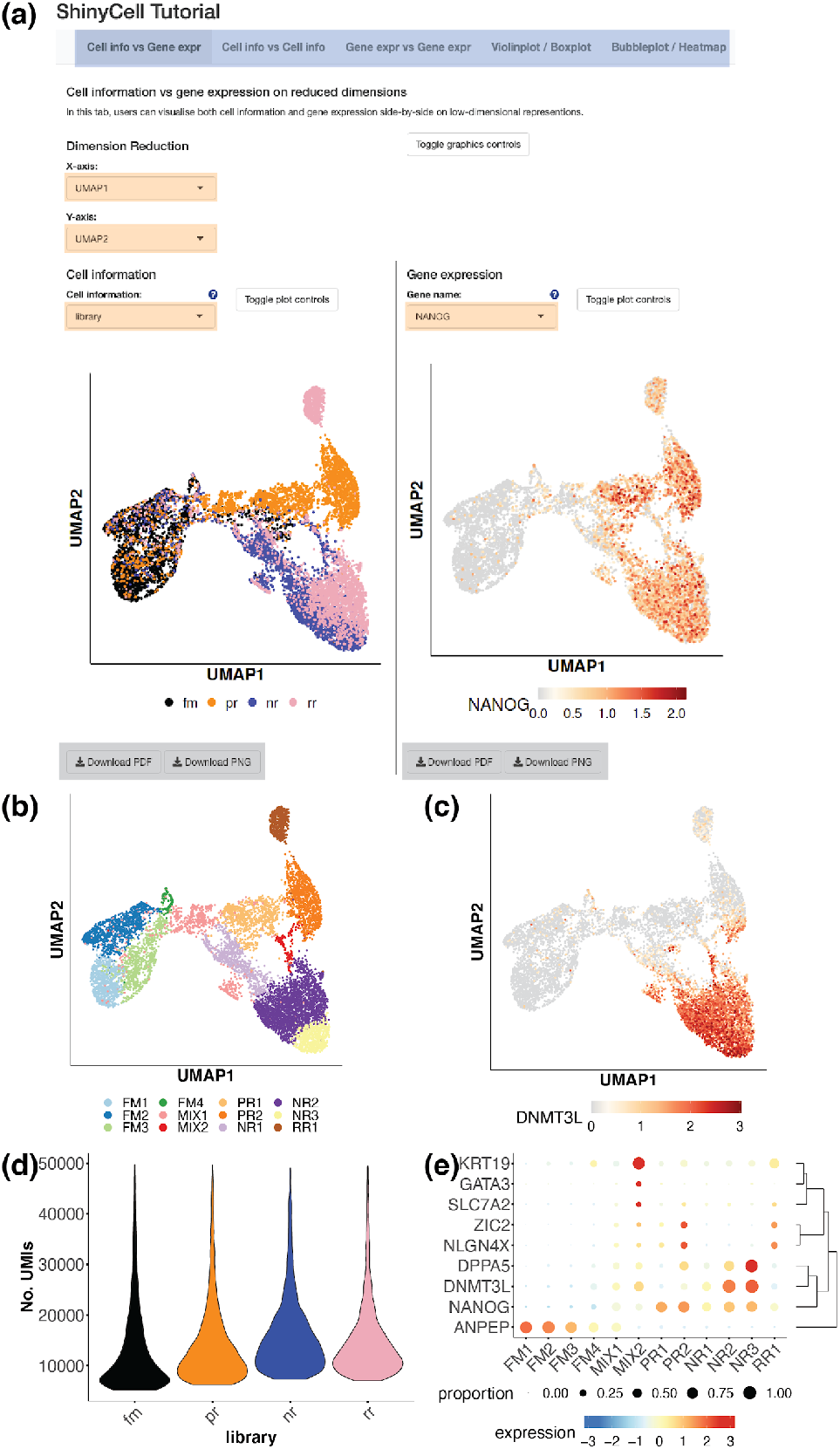
(a) An example of a ShinyCell web interface that incorporates different visualisation of single-cell data (in different tabs, highlighted in blue) that can be exported into PDF / PNG images (export buttons in grey). The ShinyCell interface allows easy comparison of gene expression with experimental parameters as well as comparing multiple features simultaneously (interactive controls highlighted in orange). Different types of visualisation can be generated from a ShinyCell web interface, including (b) cell information e.g cluster membership on low-dimensional projections, (c) gene expression on low-dimensional representations, (d) violin plots or box plots of cell information e.g. number of UMIs or gene expression and (e) bubble plots or heatmaps of the expression of multiple genes. The single-cell data is taken from a recent study of reprogramming fibroblast into primed and naive induced pluripotent stem cells (Liu *et al*., 2020).

### 2.2 Visualisation in ShinyCell interactive interface

The ShinyCell interactive web interface includes several visualisations that are common in the analysis of single-cell data. Either the cell metadata (e.g. library or cluster membership) or gene expression can be plotted on reduced dimensions (Figure 1b,c), for instance, a Uniform Manifold Approximation and Projection (McInnes *et al*., 2018) (UMAP). In the ShinyCell interface, two reduced dimension plots coloured by different information are displayed beside each other (Figure 1a). This allows users who have identified a cell population expressing a particular gene from the first plot to check the identity of the same cell population in the second plot. Alternatively, users can also check if a cell population shares similar metadata or co-expresses two genes of interest. In addition, violin plots or box plots of cell metadata e.g. the number of Unique Molecular Identifiers (UMIs) or gene expression information can be visualised (Figure 1d). Users can also plot the expression of multiple genes in the form of bubble plots where the colour and size of the bubble represent the relative expression levels and proportion of cells expressing the gene respectively (Figure 1e). Overall, the different visualisations allow users to thoroughly and quickly explore the rich single-cell data.

### 2.3 Basic usage

The ShinyCell R package takes in a pre-processed single-cell object, which can be either a Seurat or SCE object, and generates a ShinyCell configuration object containing labelling and colour palette information regarding the single-cell metadata. The ShinyCell configuration and single-cell object are then used to generate the files and code required for the Shiny app. This entire process can be easily executed using only two simple steps, shown in this example below:

~~~
#STEP 1: load a seurat object
getExampleData()
seu = readRDS(“readySeu_rset.rds”)
#STEP 2 make a ShinyCell interface
scConf = createConfig(seu)
makeShinyApp(seu, scConf, gene.mapping=TRUE,
           shiny.title=“ShinyCell Tutorial”)
~~~

The output of ShinyCell comprises (i) memory optimised files storing the single-cell dataset and (ii) two R scripts (server.R and ui.R) containing the code for the Shiny app. The Shiny app can then be opened locally via RStudio (Team and Others, 2020) or be hosted online. As the code for the Shiny app is readily available, users can add additional visualisations tailored to their single-cell analysis manually.

### 2.4 Advanced usage

The aesthetics and contents of the Shiny app can be customised by modifying the ShinyCell configuration. Users can exclude certain single-cell metadata or change the order in which the metadata are presented in the Shiny app. This is useful to organise the metadata and remove any repetitive metadata. Users can also modify the labels and 0submissiocolour palettes of categorical metadata e.g. library or cluster to match the way they are presented in the publication. Furthermore, ShinyCell automatically chooses default metadata and genes to be plotted on the Shiny app and these defaults can be altered as well. To facilitate the customisation process, we included a function that plots the legends associated with all single-cell metadata so as to provide an overview of the contents of the Shiny app. These customisations result in a more interpretable aesthetic interface making it easier to explore the data.

Another advanced use case would be the inclusion of multiple single-cell datasets into a single Shiny app. As the data files and R scripts are generated separately and in a modular manner, ShinyCell can support the inclusion of any number of single-cell datasets. For each dataset, a set of data files e.g metadata and gene expression are created with an internal dataset-specific identifier. ShinyCell then generates the Shiny app scripts, which separate the visualisation for the various datasets in different tabs.

In summary, ShinyCell allows for simple and interactive visualisation of single-cell data, which can be shared on online platforms. This will facilitate further exploration of published single-cell studies to gain additional biological insights.

## Author Contributions and Funding

O.J.L.R and J.F.O conceived and designed the ShinyCell package. J.F.O developed the code, implemented the algorithm with help from U.S.K and E.Y.C. O.J.L.R and J.F.O wrote the manuscript with input from U.S.K and E.Y.C. This work was supported by a Singapore National Research Foundation grant [NRF-CRP20-2017-0002]. Conflict of Interest: none declared.

