## Supplementary Information for "ShinyCell: Simple and sharable visualisation of single-cell gene expression data"

### Supplementary Information: ShinyCell package tutorial

`ShinyCell` is a R package that allows users to create interactive Shiny-based web applications to visualise single-cell data via (i) visualising cell information and/or gene expression on reduced dimensions e.g. UMAP, (ii) visualising the distribution of continuous cell information e.g. nUMI / module scores using violin plots / box plots and (iii) visualising the expression of multiple genes using bubbleplots / heatmap.

The package supports Seurat objects (v3.0 and above) and SingleCellExperiment objects. It is easy to use and customise settings e.g. label names and colour palettes. This readme is broken down into the following sections:

- Installation on how to install `ShinyCell`
- Quick Start Guide to rapidly deploy a shiny app with a few lines of code
- Detailed Tutorial to customise the shiny app
- Multi-dataset Tutorial to include multiple single-cell datasets into a single shiny app
- Frequently Asked Questions

### Installation

First, users can run the following code to check if the packages required by `ShinyCell` exist and install them if required:

```
reqPkg = c("data.table", "Matrix", "hdf5r", "ggplot2", "gridExtra",
           "glue", "readr", "RColorBrewer", "R.utils", "Seurat")
newPkg = reqPkg[!(reqPkg %in% installed.packages()[,"Package"])]
if(length(newPkg)){install.packages(newPkg)}
```

Furthermore, on the system where the Shiny app will be deployed, users can run the following code to check if the packages required by the Shiny app exist and install them if required:

```
reqPkg = c("shiny", "shinyhelper", "data.table", "Matrix", "hdf5r",
           "ggplot2", "gridExtra", "magrittr", "ggdendro")
newPkg = reqPkg[!(reqPkg %in% installed.packages()[,"Package"])]
if(length(newPkg)){install.packages(newPkg)}
```

`ShinyCell` can then be installed from GitHub as follows:

```
devtools::install_github("SGDDNB/ShinyCell")
```

### Quick Start Guide

In short, the `ShinyCell` package takes in an input single-cell object and generates a ShinyCell config `scConf` containing labelling and colour palette information for the single-cell metadata. The ShinyCell config and single-cell object are then used to generate the files and code required for the shiny app.

In this example, we will use single-cell data (Seurat object) containing intermediates collected during the reprogramming of human fibroblast into induced pluripotent stem cells using the RSeT media condition, taken from Liu, Ouyang, Rossello et al. Nature (2020). The Seurat object can be downloaded here.

A shiny app can then be quickly generated using the following code:

```r
library(Seurat)
library(ShinyCell)

getExampleData()                       # Download example dataset (~200 MB)
seu = readRDS("readySeu_rset.rds")
scConf = createConfig(seu)
makeShinyApp(seu, scConf, gene.mapping = TRUE,
             shiny.title = "ShinyCell Quick Start")
```

The generated shiny app can then be found in the `shinyApp/` folder (which is the default output folder). To run the app locally, use RStudio to open either `server.R` or `ui.R` in the shiny app folder and click on "Run App" in the top right corner. The shiny app can also be deployed online via online platforms e.g. shinyapps.io or be hosted via Shiny Server. The shiny app contains five tabs (highlighted in blue box), looking like this:

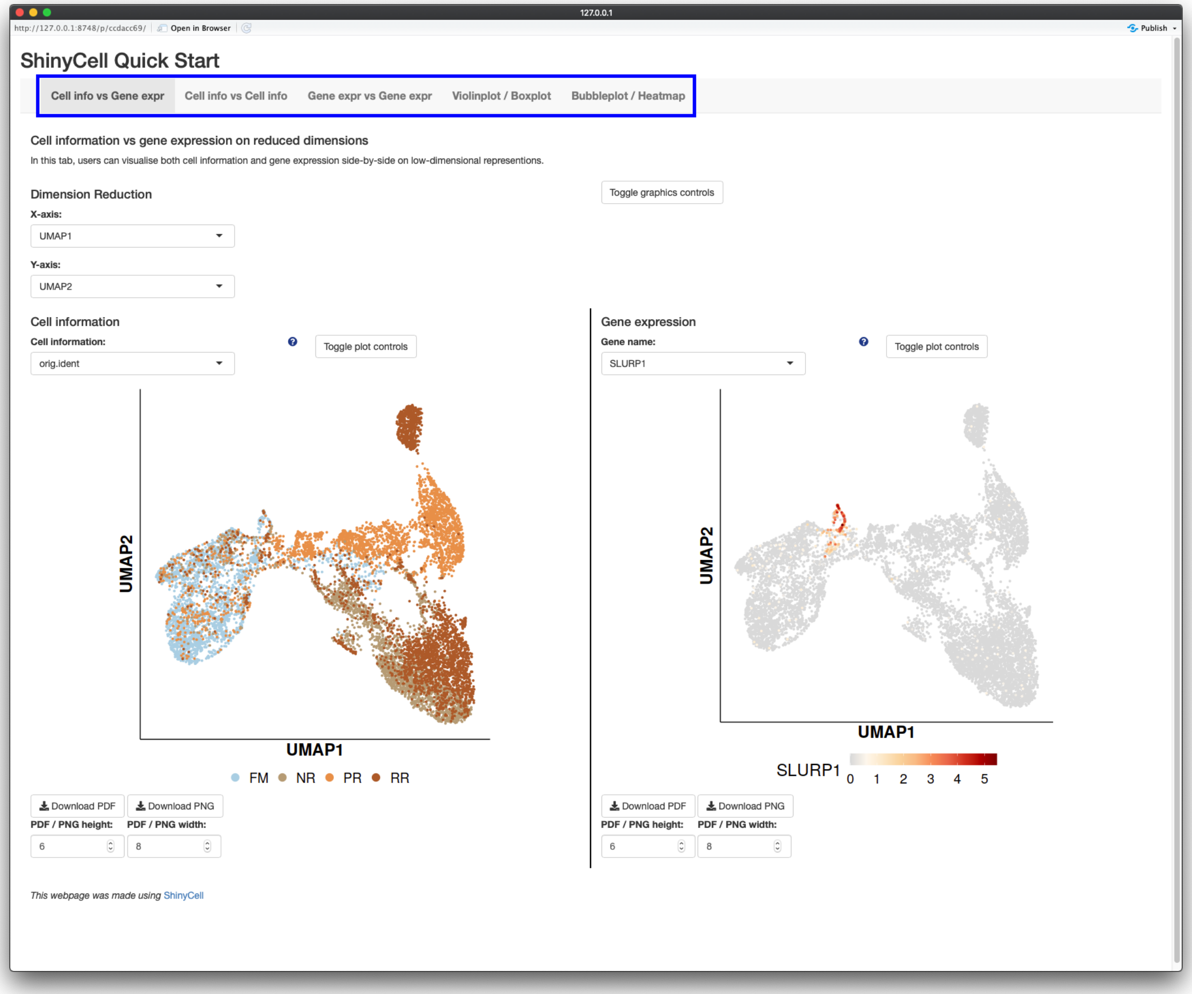


The first three tabs allows users to visualise single cells on reduced dimensions, either showing both cell information and gene expression (first tab), showing two cell information side-by-side (second tab) or showing two gene expressions side-by-side (third tab). The fourth tab allows users to visualise the distribution of continuous cell information e.g. nUMI / module scores using a violin plot or box plot and the fifth tab allows users to visualise the expression of multiple genes using a bubbleplot or heatmap.

### Detailed Tutorial

Here, we present a detailed walkthrough on how `ShinyCell` can be used to create a Shiny app from single-cell data objects, namely Seurat and SingleCellExperiment objects. In particular, we will focus on how users can customise what metadata is to be included, their labels and colour palettes.

To demonstrate, we will again use single-cell data (Seurat object) containing intermediates collected during the reprogramming of human fibroblast into induced pluripotent stem cells using the RSeT media condition, taken from Liu, Ouyang, Rossello et al. Nature (2020). The Seurat object can be downloaded here.

First, we will load the Seurat object and run `createConfig()` to create a ShinyCell configuration `scConf`. The `scConf` is a data.table containing (i) the single-cell metadata to display on the Shiny app, (ii) ordering of factors / categories for categorical metadata e.g. library / cluster and (iii) colour palette associated with each metadata. Thus, `scConf` acts as an "instruction manual" to build the Shiny app without modifying the original single-cell data.

```
library(Seurat)
library(ShinyCell)

# Create ShinyCell config
getExampleData()                       # Download example dataset (~200 MB)
seu <- readRDS("readySeu_rset.rds")
scConf = createConfig(seu)
```

To visualise the contents of the Shiny app prior to building the actual app, we can run `showLegend()` to display the legends associated with all the single-cell metadata. This allows users to visually inspect which metadata to be shown on the Shiny app. This is useful for identifying repetitive metadata and checking how factors / categories for categorical metadata will look in the eventual Shiny app. Categorical metadata and colour palettes are shown first, followed by continuous metadata which are shown collectively.

```
showLegend(scConf)
```

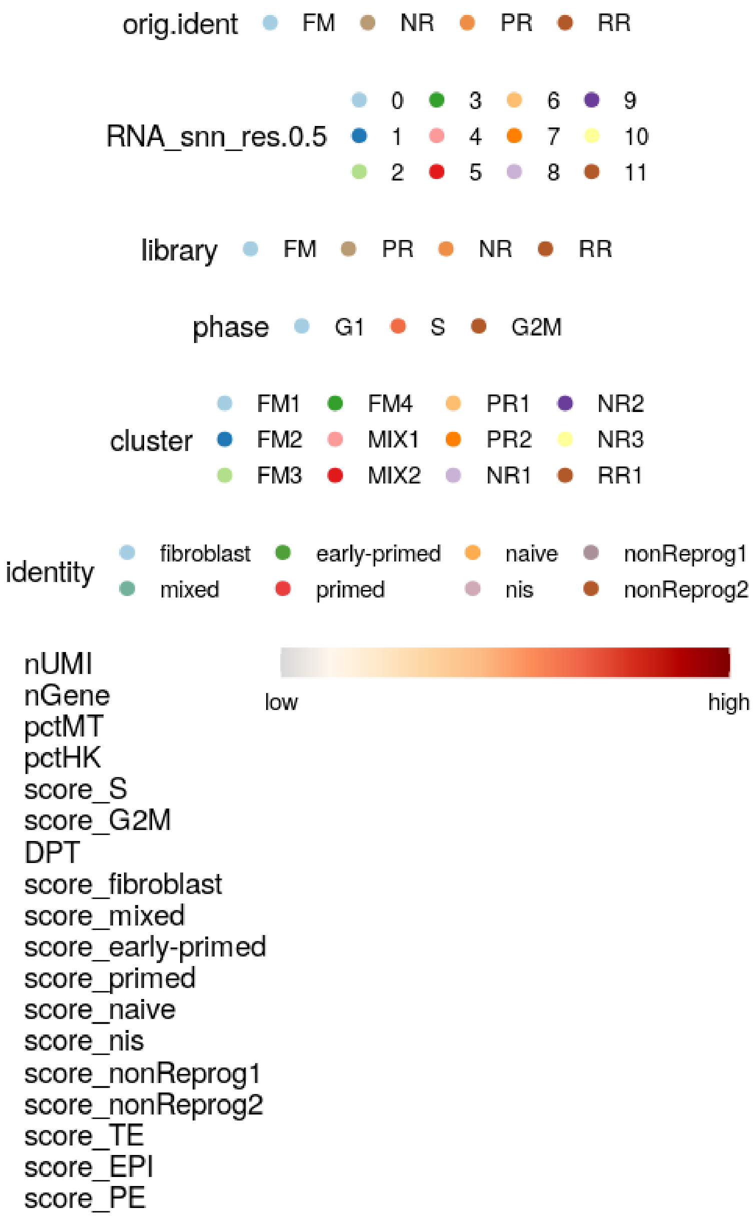


It is possible to modify `scConf` directly but this might be prone to error. Thus, we provided numerous convenience functions to modify `scConf` and ultimately the Shiny app. In this example, we note that the `orig.ident` and `library` as well as `RNA_snn_res.0.5` and `cluster` metadata are similar. To exclude metadata from the Shiny app, we can run `delMeta()`. Furthermore, we can modify how the names of metadata appear by running `modMetaName()`. In this case, we changed the names of some metadata to make them more meaningful.

By default, colours for categorical metadata are generated by interpolating colours from the "Paired" colour palette in the RColorBrewer package. To modify the colour palette, we can run `modColours()`. Here, we changed the colours for the library metadata to match that in the publication. It is also possible to modify the labels for each category via `modLabels()`. For example, we changed the labels for the library metadata from upper case to lower case. After modifying `scConf`, it is reccomended to run `showLegend()` to inspect the changes made.

```
# Delete excessive metadata and rename some metadata
scConf = delMeta(scConf, c("orig.ident", "RNA_snn_res.0.5", "phase"))
scConf = modMetaName(scConf,
                     meta.to.mod = c("nUMI", "nGene", "pctMT", "pctHK"),
                     new.name = c("No. UMIs", "No. detected genes",
                                  "% MT genes", "% HK genes"))
showLegend(scConf)

# Modify colours and labels
scConf = modColours(scConf, meta.to.mod = "library",
                    new.colours= c("black", "darkorange", "blue", "pink2"))
scConf = modLabels(scConf, meta.to.mod = "library",
                   new.labels = c("fm", "pr", "nr", "rr"))
showLegend(scConf)
```

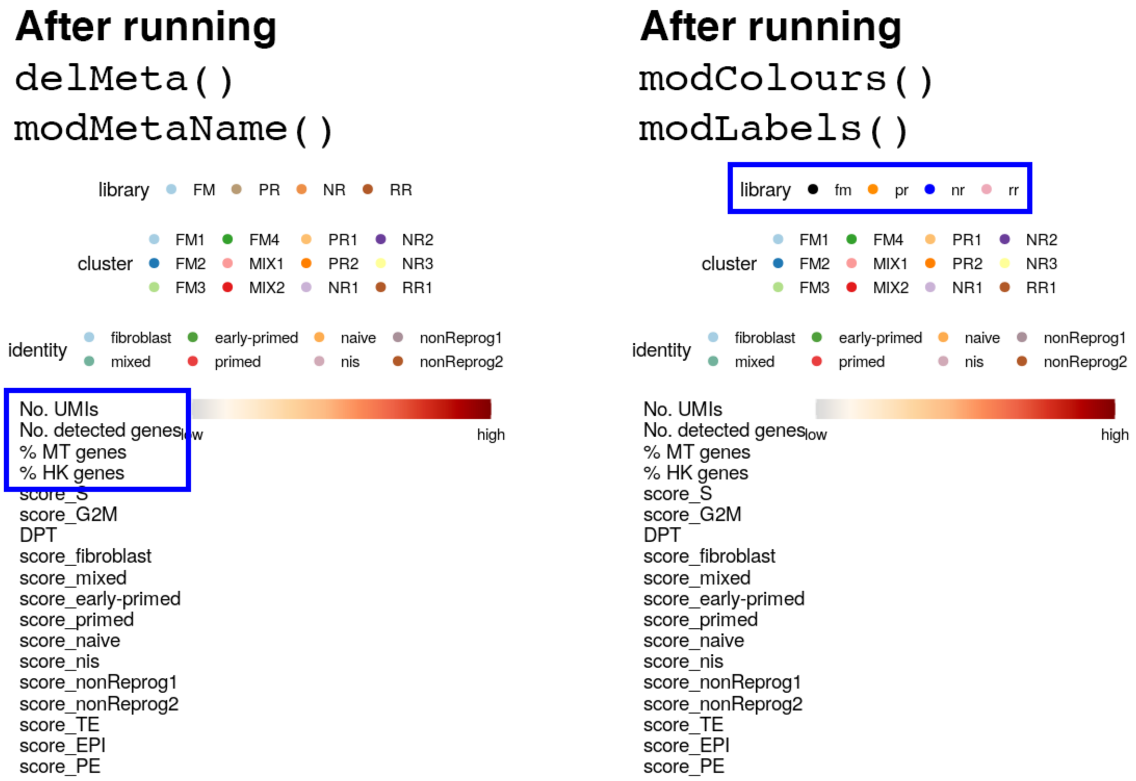


Apart from `showLegend()`, users can also run `showOrder()` to display the order in which metadata will appear in the dropdown menu when selecting which metadata to plot in the Shiny app. A table will be printed showing the actual name of the metadata in the single-cell object and the display name in the Shiny app. The metadata type (either categorical or continuous) is also provided with the number of categories "nlevels". Finally, the "default" column indicates which metadata are the primary and secondary default.

```
showOrder(scConf)
```

| | actual name | display name | type | nlevels | default |
|---|---|---|---|---|---|
| 1 | library | library | cat. | 4 | 1 |
| 2 | nUMI | No. UMIs | cont. | 0 | |
| 3 | nGene | No. detected genes | cont. | 0 | |
| 4 | pctMT | % MT genes | cont. | 0 | |
| 5 | pctHK | % HK genes | cont. | 0 | |
| 6 | score_S | score_S | cont. | 0 | |
| 7 | score_G2M | score_G2M | cont. | 0 | |
| 8 | cluster | cluster | cat. | 12 | 2 |
| 9 | identity | identity | cat. | 8 | |
| 10 | DPT | DPT | cont. | 0 | |
| 11 | score_fibroblast | score_fibroblast | cont. | 0 | |
| 12 | score_mixed | score_mixed | cont. | 0 | |
| 13 | score_early-primed | score_early-primed | cont. | 0 | |
| 14 | score_primed | score_primed | cont. | 0 | |
| 15 | score_naive | score_naive | cont. | 0 | |
| 16 | score_nis | score_nis | cont. | 0 | |
| 17 | score_nonReprog1 | score_nonReprog1 | cont. | 0 | |
| 18 | score_nonReprog2 | score_nonReprog2 | cont. | 0 | |
| 19 | score_TE | score_TE | cont. | 0 | |
| 20 | score_EPI | score_EPI | cont. | 0 | |
| 21 | score_PE | score_PE | cont. | 0 | |

Here, we introduce a few more functions that might be useful in modifying the Shiny app. Users can add metadata back via `addMeta()`. The newly added metadata (in this case, the phase metadata) is appended to the bottom of the list as shown by `showOrder()`. Next, we can reorder the order in which metadata appear in the dropdown menu in the Shiny app via `reorderMeta()`. Here, we shifted the phase metadata up the list. Finally, users can change the default metadata to plot via `modDefault()`. Again, it is reccomended to run `showOrder()` frequently to check how the metadata is changed.

```
# Add metadata back, reorder, default
scConf = addMeta(scConf, "phase", seu)
showOrder(scConf)
scConf = reorderMeta(scConf, scConf$ID[c(1:5,22,6:21)])
showOrder(scConf)
scConf = modDefault(scConf, "library", "identity")
showOrder(scConf)
```

##### addMeta()

| | actual name | display name | type | nlevels | default |
|---|---|---|---|---|---|
| 1 | library | library | cat. | 4 | 1 |
| 2 | nUMI | No. UMIs | cont. | 0 | |
| 3 | nGene | No. detected genes | cont. | 0 | |
| 4 | pctMT | % MT genes | cont. | 0 | |
| 5 | pctHK | % HK genes | cont. | 0 | |
| 6 | score_S | score_S | cont. | 0 | |
| 7 | score_G2M | score_G2M | cont. | 0 | |
| 8 | cluster | cluster | cat. | 12 | 2 |
| 9 | identity | identity | cat. | 8 | |
| 10 | DPT | DPT | cont. | 0 | |
| 11 | score_fibroblast | score_fibroblast | cont. | 0 | |
| 12 | score_mixed | score_mixed | cont. | 0 | |
| 13 | score_early-primed | score_early-primed | cont. | 0 | |
| 14 | score_primed | score_primed | cont. | 0 | |
| 15 | score_naive | score_naive | cont. | 0 | |
| 16 | score_nis | score_nis | cont. | 0 | |
| 17 | score_nonReprog1 | score_nonReprog1 | cont. | 0 | |
| 18 | score_nonReprog2 | score_nonReprog2 | cont. | 0 | |
| 19 | score_TE | score_TE | cont. | 0 | |
| 20 | score_EPI | score_EPI | cont. | 0 | |
| 21 | score_PE | score_PE | cont. | 0 | |
| 22 | phase | phase | cat. | 3 | |

##### reorderMeta()

| | actual name | display name | type | nlevels | default |
|---|---|---|---|---|---|
| 1 | library | library | cat. | 4 | 1 |
| 2 | nUMI | No. UMIs | cont. | 0 | |
| 3 | nGene | No. detected genes | cont. | 0 | |
| 4 | pctMT | % MT genes | cont. | 0 | |
| 5 | pctHK | % HK genes | cont. | 0 | |
| 6 | phase | phase | cat. | 3 | |
| 7 | score_S | score_S | cont. | 0 | |
| 8 | score_G2M | score_G2M | cont. | 0 | |
| 9 | cluster | cluster | cat. | 12 | 2 |
| 10 | identity | identity | cat. | 8 | |
| 11 | DPT | DPT | cont. | 0 | |
| 12 | score_fibroblast | score_fibroblast | cont. | 0 | |
| 13 | score_mixed | score_mixed | cont. | 0 | |
| 14 | score_early-primed | score_early-primed | cont. | 0 | |
| 15 | score_primed | score_primed | cont. | 0 | |
| 16 | score_naive | score_naive | cont. | 0 | |
| 17 | score_nis | score_nis | cont. | 0 | |
| 18 | score_nonReprog1 | score_nonReprog1 | cont. | 0 | |
| 19 | score_nonReprog2 | score_nonReprog2 | cont. | 0 | |
| 20 | score_TE | score_TE | cont. | 0 | |
| 21 | score_EPI | score_EPI | cont. | 0 | |
| 22 | score_PE | score_PE | cont. | 0 | |

##### modDefault()

| | actual name | display name | type | nlevels | default |
|---|---|---|---|---|---|
| 1 | library | library | cat. | 4 | 1 |
| 2 | nUMI | No. UMIs | cont. | 0 | |
| 3 | nGene | No. detected genes | cont. | 0 | |
| 4 | pctMT | % MT genes | cont. | 0 | |
| 5 | pctHK | % HK genes | cont. | 0 | |
| 6 | phase | phase | cat. | 3 | |
| 7 | score_S | score_S | cont. | 0 | |
| 8 | score_G2M | score_G2M | cont. | 0 | |
| 9 | cluster | cluster | cat. | 12 | |
| 10 | identity | identity | cat. | 8 | 2 |
| 11 | DPT | DPT | cont. | 0 | |
| 12 | score_fibroblast | score_fibroblast | cont. | 0 | |
| 13 | score_mixed | score_mixed | cont. | 0 | |
| 14 | score_early-primed | score_early-primed | cont. | 0 | |
| 15 | score_primed | score_primed | cont. | 0 | |
| 16 | score_naive | score_naive | cont. | 0 | |
| 17 | score_nis | score_nis | cont. | 0 | |
| 18 | score_nonReprog1 | score_nonReprog1 | cont. | 0 | |
| 19 | score_nonReprog2 | score_nonReprog2 | cont. | 0 | |
| 20 | score_TE | score_TE | cont. | 0 | |
| 21 | score_EPI | score_EPI | cont. | 0 | |
| 22 | score_PE | score_PE | cont. | 0 | |

After modifying `scConf` to one's satisfaction, we are almost ready to build the Shiny app. Prior to building the Shiny app, users can run `checkConfig()` to check if the `scConf` is ready. This is especially useful if users have manually modified the `scConf`. Users can also add a footnote to the Shiny app and one potential use is to include the reference for the dataset. Here, we provide an example of including the citation as the Shiny app footnote.

```
# Build shiny app
checkConfig(scConf, seu)
footnote = paste0(
  'strong("Reference: "), "Liu X., Ouyang J.F., Rossello F.J. et al. ",',
  'em("Nature "), strong("586,"), "101-107 (2020) ",',
  'a("doi:10.1038/s41586-020-2734-6",',
  'href = "https://www.nature.com/articles/s41586-020-2734-6",',
  'target="_blank"), style = "font-size: 125%;"'
)
```

Now, we can build the shiny app! A few more things need to be specified here. In this example, the Seurat object uses Ensembl IDs and we would like to convert them to more user-friendly gene symbols in the Shiny app. `ShinyCell` can do this conversion (for human and mouse datasets) conveniently by specifying `gene.mapping = TRUE`. If your dataset is already in gene symbols, you can leave out this argument to not perform the conversion. Furthermore, `ShinyCell` uses the "RNA" assay and "data" slot in Seurat objects as the gene expression data. If you have performed any data integration and would like to use the integrated data instead, please specify `gex.assay = "integrated`. Also, default genes to plot can be specified where `default.gene1` and `default.gene2` corresponds to the default genes when plotting gene expression on reduced dimensions while `default.multigene` contains the default set of multiple genes when plotting bubbleplots or heatmaps. If unspecified, `ShinyCell` will automatically select some genes present in the dataset as default genes.

```
makeShinyApp(seu, scConf, gene.mapping = TRUE,
             gex.assay = "RNA", gex.slot = "data",
             shiny.title = "ShinyCell Tutorial",
             shiny.dir = "shinyApp/", shiny.footnotes = footnote,
             default.gene1 = "NANOG", default.gene2 = "DNMT3L",
             default.multigene = c("ANPEP","NANOG","ZIC2","NLGN4X","DNMT3L",
                                   "DPPA5","SLC7A2","GATA3","KRT19"))
```

Under the hood, `makeShinyApp()` does two things: generate (i) the data files required for the Shiny app and (ii) the code files, namely `server.R` and `ui.R`. The generated files can be found in the `shinyApp/` folder. To run the app locally, use RStudio to open either `server.R` or `ui.R` in the shiny app folder and click on "Run

App” in the top right corner. The shiny app can also be deployed via online platforms e.g. shinyapps.io or hosted via Shiny Server. The shiny app look like this, containing five tabs. Cell information and gene expression are plotted on UMAP in the first tab while two different cell information / gene expression are plotted on UMAP in the second / third tab respectively. Violin plot or box plot of cell information or gene expression distribution can be found in the fourth tab. Lastly, a bubbleplot or heatmap can be generated in the fifth tab.

With the Shiny app, users can interactively explore their single-cell data, varying the cell information / gene expression to plot. Furthermore, these plots can be exported into PDF / PNG for presentations / publications. Users can also click on the “Toggle graphics controls” or “Toggle plot controls” to fine-tune certain aspects of the plots e.g. point size.

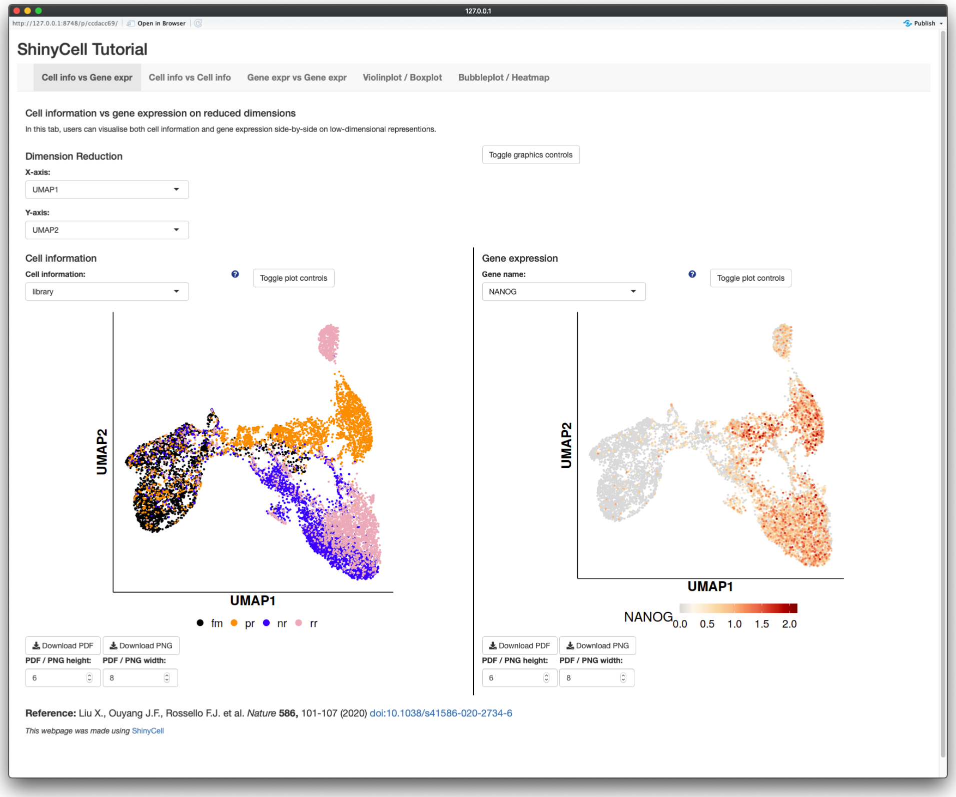

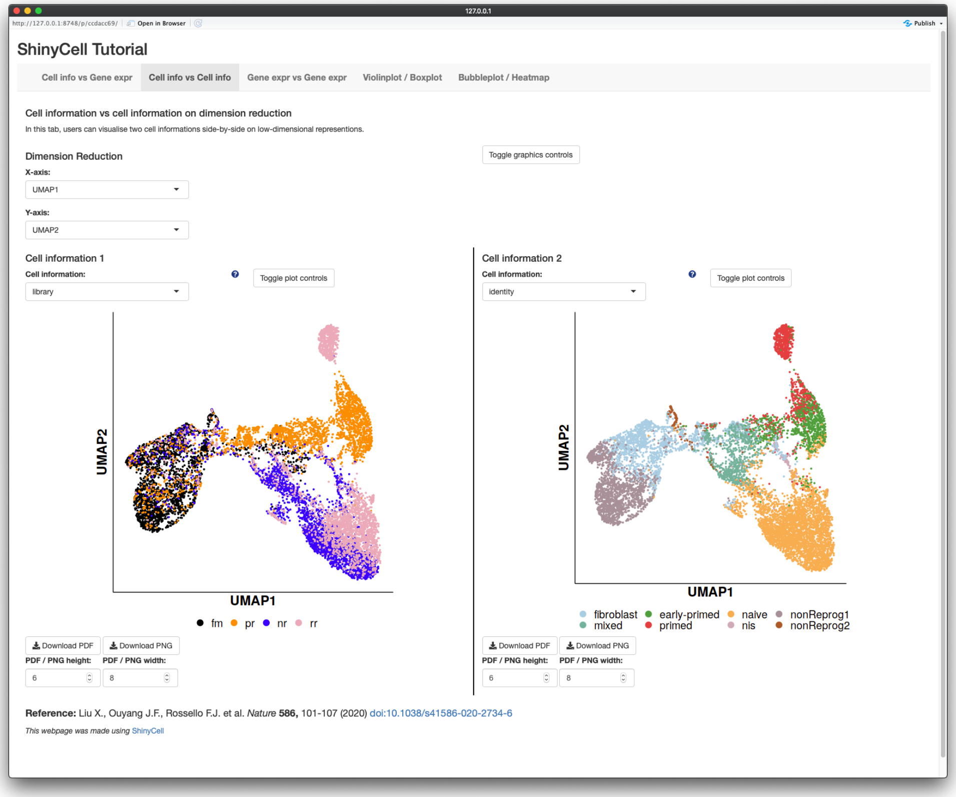
ShinyCell Tutorial
Cell info vs Gene expr
Cell info vs Cell info
Gene expr vs Gene expr
Violinplot / Boxplot
Bubbleplot / Heatmap
Cell information vs cell information on dimension reduction
In this tab, users can visualise two cell informations side-by-side on low-dimensional represention.
Dimension Reduction
Toggle graphics controls
X-axis:
UMAP1
Y-axis:
UMAP2
Cell information 1
Cell information:
library
Toggle plot controls
Cell information 2
Cell information:
identity
Toggle plot controls
UMAP2
UMAP1
fm
pr
nr
rr
fibroblast
mixed
early-primed
primed
naive
nis
nonReprog1
nonReprog2
Download PDF
Download PNG
PDF / PNG height:
PDF / PNG width:
Reference: Liu X., Ouyang J.F., Rossello F.J. et al. Nature 586, 101-107 (2020) doi:10.1038/s41586-020-2734-6
This webpage was made using ShinyCell

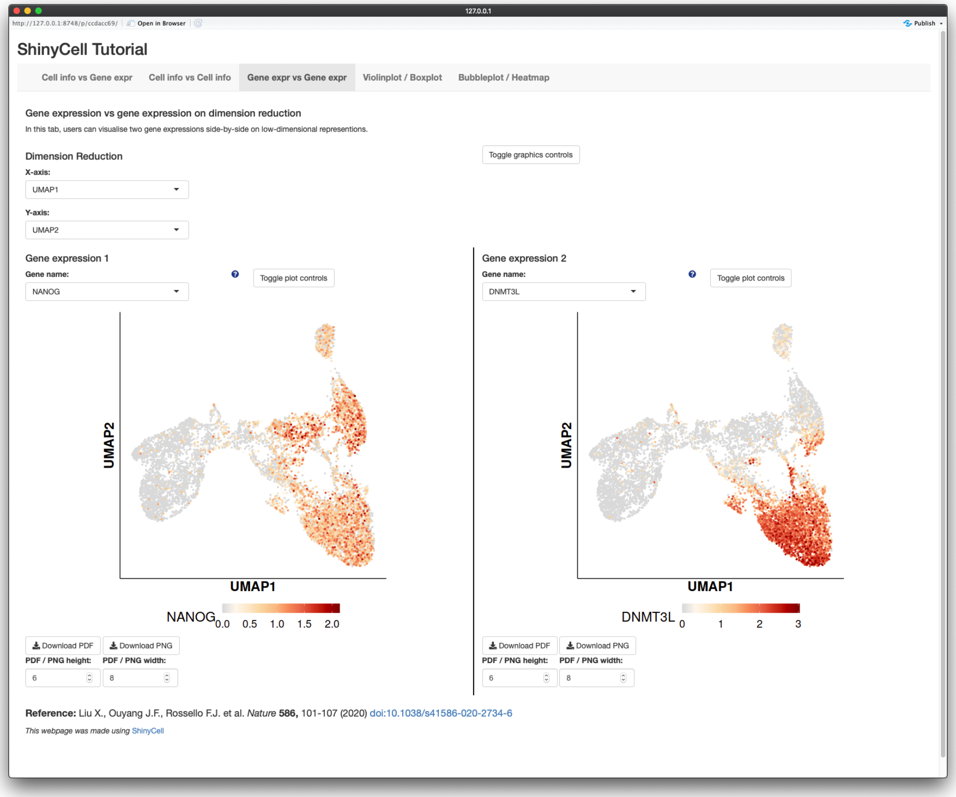
ShinyCell Tutorial
Cell info vs Gene expr
Cell info vs Cell info
Gene expr vs Gene expr
Violinplot / Boxplot
Bubbleplot / Heatmap
Gene expression vs gene expression on dimension reduction
In this tab, users can visualise two gene expressions side-by-side on low-dimensional representions.
Dimension Reduction
Toggle graphics controls
X-axis:
UMAP1
Y-axis:
UMAP2
Gene expression 1
Gene name:
NANOG
Toggle plot controls
Gene expression 2
Gene name:
DNMT3L
Toggle plot controls
UMAP2
UMAP1
NANOG
0.0
0.5
1.0
1.5
2.0
DNMT3L
0
1
2
3
Download PDF
Download PNG
PDF / PNG height:
PDF / PNG width:
Reference: Liu X., Ouyang J.F., Rossello F.J. et al. Nature 586, 101-107 (2020) doi:10.1038/s41586-020-2734-6
This webpage was made using ShinyCell

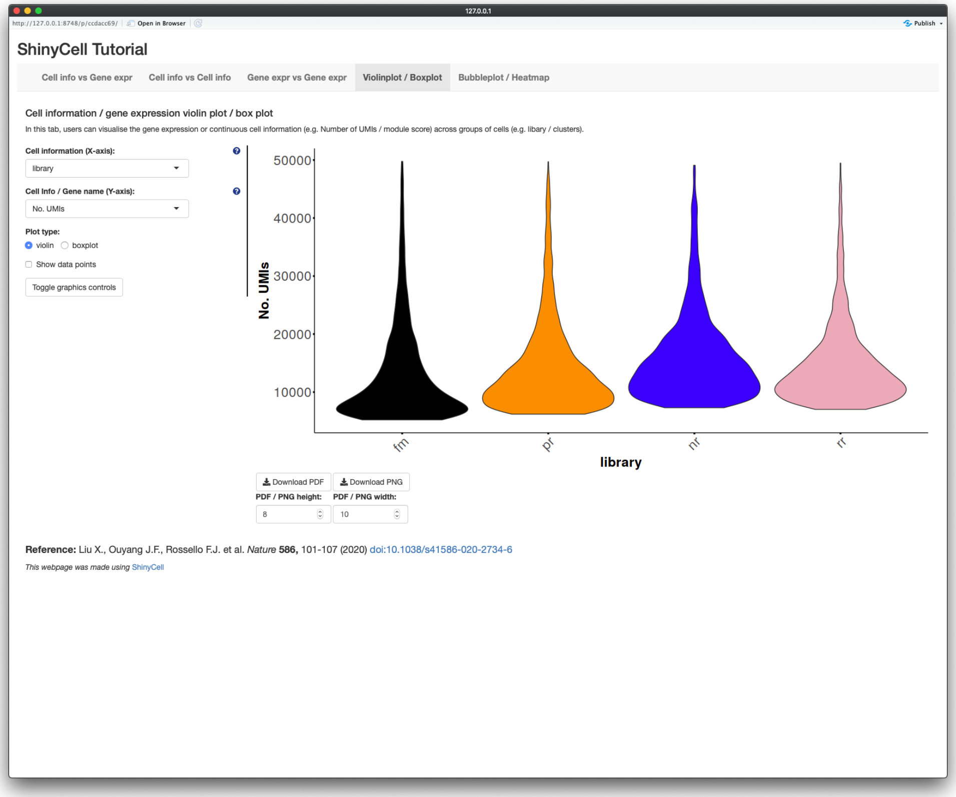
ShinyCell Tutorial
Cell info vs Gene expr
Cell info vs Cell info
Gene expr vs Gene expr
Violinplot / Boxplot
Bubbleplot / Heatmap
Cell information / gene expression violin plot / box plot
In this tab, users can visualise the gene expression or continuous cell information (e.g. Number of UMIs / module score) across groups of cells (e.g. libary / clusters).
Cell information (X-axis):
library
Cell Info / Gene name (Y-axis):
No. UMIs
Plot type:
violin
boxplot
Show data points
Toggle graphics controls
No. UMIs
library
fm
pr
nr
rr
Download PDF
Download PNG
PDF / PNG height:
PDF / PNG width:
Reference: Liu X., Ouyang J.F., Rossello F.J. et al. Nature 586, 101-107 (2020) doi:10.1038/s41586-020-2734-6
This webpage was made using ShinyCell

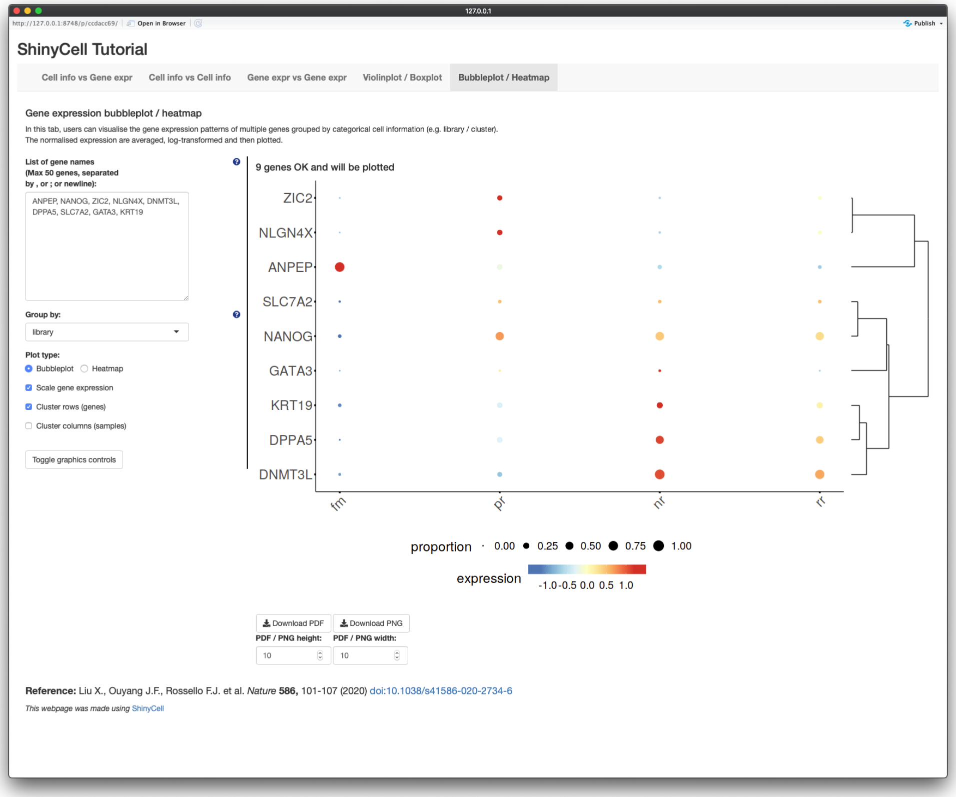
ShinyCell Tutorial
Cell info vs Gene expr
Cell info vs Cell info
Gene expr vs Gene expr
Violinplot / Boxplot
Bubbleplot / Heatmap
Gene expression bubbleplot / heatmap
In this tab, users can visualise the gene expression patterns of multiple genes grouped by categorical cell information (e.g. library / cluster).
The normalised expression are averaged, log-transformed and then plotted.
List of gene names (Max 50 genes, separated by , or ; or newline):
ANPEP, NANOG, ZIC2, NLGN4X, DNMT3L, DPPA5, SLC7A2, GATA3, KRT19
Group by:
library
Plot type:
Bubbleplot
Heatmap
Scale gene expression
Cluster rows (genes)
Cluster columns (samples)
Toggle graphics controls
9 genes OK and will be plotted
ZIC2
NLGN4X
ANPEP
SLC7A2
NANOG
GATA3
KRT19
DPPA5
DNMT3L
fm
pr
nr
rr
proportion
expression
Download PDF
Download PNG
PDF / PNG height:
PDF / PNG width:
Reference: Liu X., Ouyang J.F., Rossello F.J. et al. Nature 586, 101-107 (2020) doi:10.1038/s41586-020-2734-6
This webpage was made using ShinyCell

### Multi-dataset Tutorial

Users might want to include multiple single-cell datasets into a single Shiny app and `ShinyCell` provides this functionality. We will demonstrate how to create a Shiny app containing two single-cell datasets. For the first example dataset, we will again use scRNA-seq data (Seurat object) containing intermediates collected during the reprogramming of human fibroblast into induced pluripotent stem cells using the RSeT media condition, which can be downloaded here. For the second example dataset, we will use scRNA-seq of day 21 reprogramming intermediates from the same publication, which can be downloaded here. After downloading the data, we will begin by loading the required libraries.

```
library(Seurat)
library(ShinyCell)
getExampleData("multi")       # Download multiple example datasets (~400 MB)
```

To create a multi-dataset Shiny app, we need to configure the settings for each dataset separately. We will do so for the first dataset as follows. A ShinyCell configuration `scConf1` is created, followed by modifying various aspects of the Shiny app e.g. removing excessive metadata, modifying the display names of metadata and modifying the colour palettes. For a more detailed explanation on how to customise the shiny app, refer to the Detailed Tutorial. We then run `makeShinyFiles()` to generate the files related to the first dataset. Notice that we specified `shiny.prefix = "sc1"` and this prefix is used to identify that the files contain single-cell data related to the first dataset. The remaining arguments are the same as explained in the Detailed Tutorial.

```
seu <- readRDS("readySeu_rset.rds")
scConf1 = createConfig(seu)
scConf1 = delMeta(scConf1, c("orig.ident", "RNA_snn_res.0.5"))
scConf1 = modMetaName(scConf1, meta.to.mod = c("nUMI", "nGene", "pctMT", "pctHK"),
                      new.name = c("No. UMIs", "No. detected genes",
                                   "% MT genes", "% HK genes"))
scConf1 = modColours(scConf1, meta.to.mod = "library",
                     new.colours= c("black", "darkorange", "blue", "pink2"))
makeShinyFiles(seu, scConf1, gex.assay = "RNA", gex.slot = "data",
               gene.mapping = TRUE, shiny.prefix = "sc1",
               shiny.dir = "shinyAppMulti/",
               default.gene1 = "NANOG", default.gene2 = "DNMT3L",
               default.multigene = c("ANPEP","NANOG","ZIC2","NLGN4X","DNMT3L",
                                     "DPPA5","SLC7A2","GATA3","KRT19"),
               default.dimred = c("UMAP_1", "UMAP_2"))
```

We then repeat the same procedure for the second dataset to generate the files required for the Shiny app. Notice that we used a different prefix here `shiny.prefix = "sc2"`.

```
seu <- readRDS("readySeu_d21i.rds")
scConf2 = createConfig(seu)
scConf2 = delMeta(scConf2, c("orig.ident", "RNA_snn_res.0.5"))
scConf2 = modMetaName(scConf2, meta.to.mod = c("nUMI", "nGene", "pctMT", "pctHK"),
                      new.name = c("No. UMIs", "No. detected genes",
                                   "% MT genes", "% HK genes"))
scConf2 = modColours(scConf2, meta.to.mod = "library",
                     new.colours= c("black", "blue", "purple"))
makeShinyFiles(seu, scConf2, gex.assay = "RNA", gex.slot = "data",
               gene.mapping = TRUE, shiny.prefix = "sc2",
               shiny.dir = "shinyAppMulti/",
               default.gene1 = "GATA3", default.gene2 = "DNMT3L",
               default.multigene = c("ANPEP","NANOG","ZIC2","NLGN4X","DNMT3L",
                                     "DPPA5","SLC7A2","GATA3","KRT19"),
```

```
                default.dimred = c("UMAP_1", "UMAP_2"))
```

We can the proceed to the final part where we generate the code for the Shiny app using the `makeShinyCodesMulti()` function. To specify that two datasets will be included in this Shiny app, we input the prefixes of the two datasets `shiny.prefix = c("sc1", "sc2")`. Also, users need to specify section headers for each dataset via the `shiny.headers` argument. The remaining arguments are the same as explained in the Detailed Tutorial.

```
footnote = paste0(
  'strong("Reference: "), "Liu X., Ouyang J.F., Rossello F.J. et al. ",',
  'em("Nature "), strong("586,"), "101-107 (2020) ",',
  'a("doi:10.1038/s41586-020-2734-6",',
  'href = "https://www.nature.com/articles/s41586-020-2734-6",',
  'target="_blank"), style = "font-size: 125%;"'
)
makeShinyCodesMulti(
  shiny.title = "Multi-dataset Tutorial", shiny.footnotes = footnote,
  shiny.prefix = c("sc1", "sc2"),
  shiny.headers = c("RSeT reprogramming", "Day 21 intermediates"),
  shiny.dir = "shinyAppMulti/")
```

Now, we have both the data and code for the Shiny app and we can run the Shiny app. Each dataset can be found in their corresponding tabs and clicking on the tab creates a dropdown to change the type of plot to display on the Shiny app. This tutorial can be easily expanded to include three or more datasets. Users simply have to create the corresponding data files for each dataset and finally generate the code for the Shiny app.

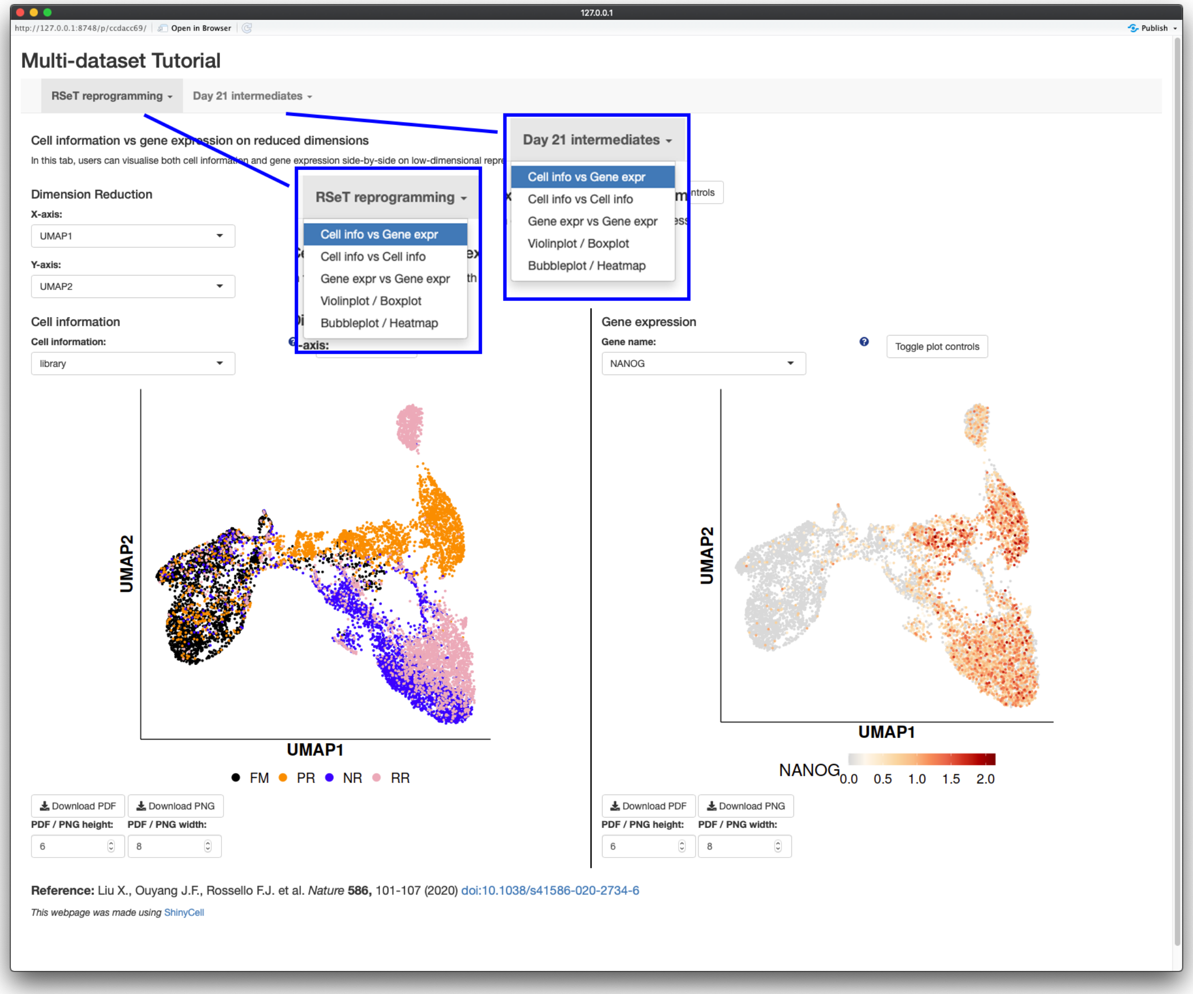


### Frequently Asked Questions

- Q: How much memory / storage space does `ShinyCell` and the Shiny app consume?
  - A: The Shiny app itself consumes very little memory and is meant to be a heavy-duty app where multiple users can access the app simultaneously. Unlike typical R objects, the entire gene expression matrix is stored on disk and *not on memory* via the hdf5 file system. Also, the hdf5 file system offers superior file compression and takes up less storage space than native R file formats such as rds / Rdata files.
  - A: It should be noted that a large amount of memory is required when *building* the Shiny app. This is because the whole Seurat / SCE object has to be loaded onto memory and additional memory is required to generate the required files. From experience, a typical laptop with 8GB RAM can handle datasets around 30k cells while 16GB RAM machines can handle around 60k-70k cells.
- Q: I have generated additional dimension reductions (e.g. force-directed layout / MDS / monocle2 etc.) and would like to include them into the Shiny app. How do I do that?
  - A: `ShinyCell` automatically retrieves dimension reduction information from the Seurat or SCE object. Thus, the additional dimension reductions have to be added into the Seurat or SCE object before running `ShinyCell`.

  - For Seurat objects, users can refer "Storing a custom dimensional reduction calculation" in https://satijalab.org/seurat/v3.0/dim_reduction_vignette.html
  - For SCE objects, users can refer to https://bioconductor.org/packages/devel/bioc/vignettes/SingleCellExperiment/inst/doc/intro.html#3_Adding_low-dimensional_representations
- Q: I have both RNA and integrated data in my Seurat object. How do I specify which gene expression assay to plot in the Shiny app?
  - A: Only one gene expression assay can be visualised per dataset. To specify the assay, use the `gex.assay = "integrated` argument in the `makeShinyApp()` or `makeShinyFiles()` functions. If users want to visualise both gene expression, they have to treat each assay as an individual dataset and include multiple datasets into a single shiny app, following the Multi-dataset Tutorial
